# The cytokine receptor DR3 identifies and activates thymic NKT17 cells

**DOI:** 10.1101/2021.09.27.461992

**Authors:** Shunqun Luo, Nurcin Liman, Assiatu Crossman, Jung-Hyun Park

**Author notes:** contributed equally (Shunqun Luo, Nurcin Liman). **Address to correspondence to:** Jung-Hyun Park, Experimental Immunology Branch, Center for Cancer Research, NCI, NIH, Building 10, Room 5B17, 10 Center Dr., Bethesda, MD 20892.

## Abstract

Invariant natural killer T (*i*NKT) cells are thymus-generated T cells with innate-like characteristics and effector function. Several functionally distinct *i*NKT subsets have been identified, but NKT17 is the only *i*NKT subset that produces the proinflammatory cytokine IL-17. NKT17 cells are generated in the thymus and then exported into the periphery to populate lymphoid organs and barrier tissues, such as the lung, to provide critical support in host defense. However, the molecular mechanisms that drive the thymic development and subset-specific activation of NKT17 cells remain mostly unknown. Here, we identify the cytokine receptor DR3, a member of the TNF receptor superfamily, being selectively expressed on NKT17 cells but absent on all other thymic *i*NKT subsets. We further demonstrate that DR3 ligation leads to the *in vivo* activation of thymic NKT17 cells and provides *in vitro* costimulatory effects upon α-GalCer-stimulation. Thus, our study reports the identification of a specific surface marker for thymic NKT17 cells that selectively triggers their activation both *in vivo* and *in vitro*. These findings provide new insights for deciphering the role and function of IL-17-producing NKT17 cells and for understanding the development and activation mechanisms of *i*NKT cells in general.

## Introduction

*i*NKT cells are thymus-derived effector T cells expressing a semi-invariant Vα14-Jα18 T cell receptor (TCR) that equips them with the ability to recognize microbial glycolipids in the context of the nonclassical MHC-I molecule CD1d. Unlike conventional αβ T cells, *i*NKT cells possess the innate ability to express effector molecules and proinflammatory cytokines prior to their exposure to antigens. While *i*NKT cells are few in their number and limited in their TCR repertoire, *i*NKT cells play critical roles in immunosurveillance, inflammation, and host defense (*Bendelac et al., 2007; Crosby et al., 2018*). There are several subsets of *i*NKT cells, among which three major populations, *i.e*., NKT1, NKT2, and NKT17, have been identified (*Lee et al., 2013*). In particular, NKT17 cells are noted for their ability to produce the proinflammatory cytokine IL-17 and to express the transcription factor RORγt (*Lee et al., 2013*). NKT17 cells can be identified by their distinct expression of the cell surface marker CD138 (Syndecan-1) (*Dai et al., 2015*). However, the role of CD138 in NKT17 cell biology remains mostly unclear (*Dai et al., 2015*)[Please add: Luo S., 2021, JCI Insight]. Because NKT17 cells are the major producers of IL-17 in the thymus and in barrier tissues, such as the lung and skin (*Tsagaratou, 2019*), there is a keen interest in delineating the developmental requirements and activation mechanism of NKT17 cells. Here, we report the surprising finding that the TNF receptor superfamily member Death Receptor-3 (DR3) is highly and specifically expressed on thymic NKT17 cells, and that the stimulation of DR3 using agonistic anti-DR3 antibodies leads to the activation of NKT17 cells in the thymus, unveiling a new layer of control in NKT17 cell biology.

## Results and Discussion

### The cytokine receptor DR3 is specifically expressed on thymic NKT17 cells

We embarked on this study to uncover new regulatory mechanisms and effector functions that are specifically associated with individual *i*NKT subsets, and particularly with NKT17 cells. While CD138 is a specific marker for NKT17 cells, CD138 is not required for their generation or effector function (*Dai et al., 2015; Luo et al., 2021*) Thus, functional markers for NKT17 cells are currently not available. Because cytokines play critical roles in the generation and survival of *i*NKT cells (*Bendelac et al., 2007; Crosby et al., 2018*), we screened a panel of cytokine receptors for their *i*NKT subset-specific expression, and here we identified the TNF receptor superfamily member 25 (TNFRS25), also known as DR3 (*Meylan et al., 2011*), being highly expressed on thymic NKT17 cells (**Figure 1A and 1B**). As expected, DR3 expression correlated closely with CD138 expression in thymic *i*NKT cells of both C57BL/6 and BALB/c mice (**Figure 1A, 1B; Figure 1–figure supplement 1**). On the other hand, DR3 expression was independent of CD138 because DR3 was still abundantly and specifically found on NKT17 cells of CD138-deficient (*Sdc1*^−/−^) BALB/c mice (**Fig. 1C**). Furthermore, the forced expression of RORγt (*Ligons et al., 2018*), the master transcription factor of NKT17 cell development and function (*Tsagaratou, 2019*), dramatically increased both the frequency and number of DR3-expressing *i*NKT cells (**Figure 1D**). These results suggested that DR3 expression is controlled downstream of RORγt so that all DR3^+^ thymic *i*NKT cells of RORγt^Tg^ mice also expressed CD138 (**Figure 1D; Figure 1–figure supplement 2**).

**Figure 1.**
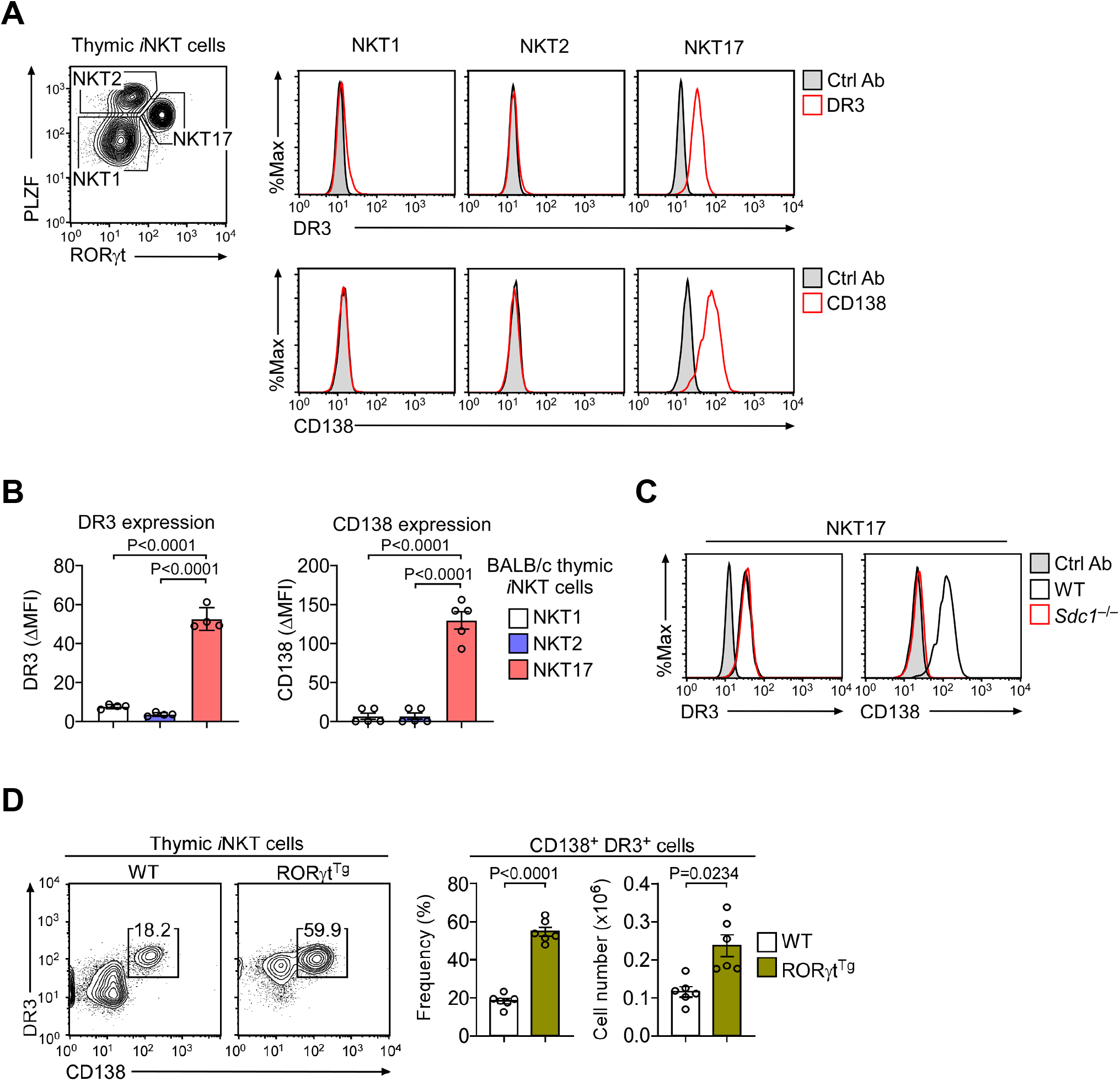
DR3 is specifically expressed on thymic NKT17 cells. **A.** *i*NKT subsets were identified among the thymocytes of BALB/c mice by staining for the intracellular proteins RORγt and PLZF and measuring the subset-specific expression of DR3 and CD138. The data are representative of 2 independent experiments with a total 4 BALB/c mice. **B.** Quantification of DR3 and CD138 expression (ΔMFI) in individual *i*NKT subsets. The data show the summary of 2 independent experiments with a total of at least 4 BALB/c mice. Statistical significance was determined by unpaired two-tailed Student’s *t*-tests. **C.** DR3 and CD138 expression on thymic NKT17 cells of CD138-deficient (*Sdc1*^−/−^) and littermate control (WT) BALB/c mice. The data are representative of 2 independent experiments with a total 4 *Sdc1*^−/−^ and 4 WT BALB/c mice. **D.** Thymic *i*NKT cells of RORγt^Tg^ and littermate control (WT) BALB/c thymocytes were assessed for surface DR3 and CD138 expression. The contour plot is representative (left), and the bar graphs (right) are a summary of data from three independent experiments with a total of 6 RORγt^Tg^ and 6 WT mice. Statistical significance was determined by paired two-tailed Student’s *t*-tests. The following figure supplements are available for Figure 1: **Figure supplement 1**. DR3 expression on thymic NKT17 cells of C57BL/6 mice. **Figure supplement 2**. Thymic CD138^+^ *i*NKT cells in WT and RORγt^Tg^ BALB/c mice.

DR3 is the receptor for the cytokine TNF-like ligand 1A (TL1A) (*Valatas et al., 2019*). Consistent with the notion that DR3 is highly expressed on Foxp3^+^ Treg cells (*Nishikii et al., 2016*), the stimulation with TL1A or the ligation of DR3 with agonistic anti-DR3 antibodies triggers the activation of Foxp3^+^ Treg cells (*Nishikii et al., 2016*). Because we found NKT17 cells to express DR3, we thus asked whether DR3 ligation would also activate thymic NKT17 cells. To this end, we injected BALB/c mice with agonistic anti-DR3 antibodies and assessed their effect on thymic *i*NKT cells. Of note, we utilized BALB/c mice that were engineered to express *Foxp3-GFP* reporter proteins (Foxp3-DTR/EGFP mice) (*Kim et al., 2007*), which allowed us to verify the *in vivo* effect of anti-DR3 injection. Indeed, assessing GFP-expressing CD4 T cells confirmed that DR3 ligation induced the expansion of Foxp3^+^ Treg cells (**Figure 2A**). Curiously, while both the frequency and number of Foxp3^+^ cells were significantly increased in DR3-injected mice, at the same time, the frequency and number of thymic NKT17 cells were dramatically diminished (**Figure 2B**). Thus, DR3 ligation clearly affected NKT17 cells, but DR3 activation appeared to be detrimental instead of stimulatory for thymic NKT17 cells.

**Figure 2.**
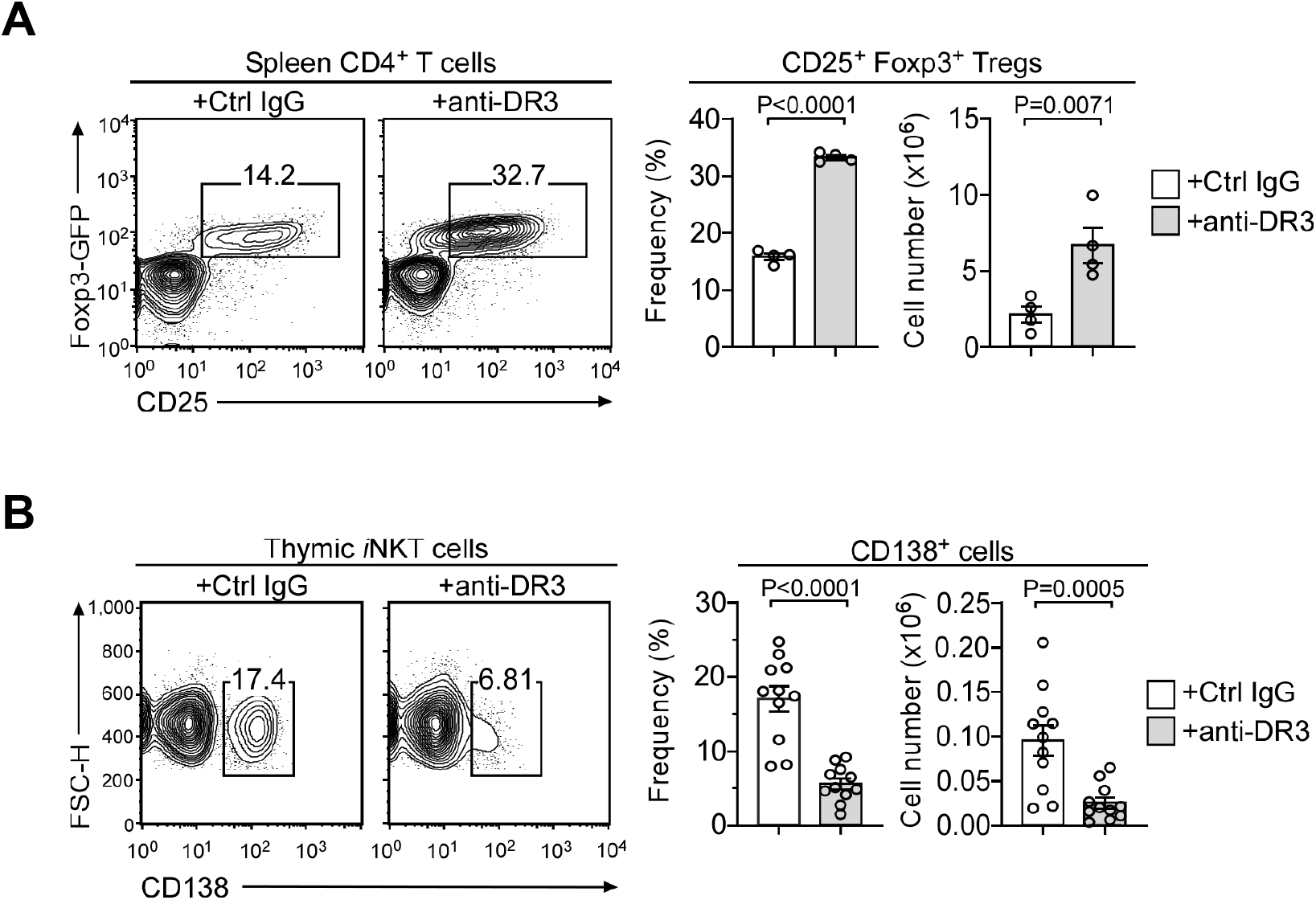
*In vivo* effects of DR3 ligation on Foxp3^+^ Treg and CD138^+^ NKT17 cells. **A.** Contour plots show *Foxp3-GFP* versus CD25 expression of spleen CD4^+^ T cells (left), and bar graphs show the frequencies and cell numbers of splenic CD25^+^Foxp3^+^ Treg cells (right), after 1 week of injection with anti-DR3 or isotype control antibodies into BALB/c Foxp3-GFP reporter mice. The results are summarized from 4 independent experiments with a total of 4 mice injected with anti-DR3 and 4 mice injected with isotype control. Statistical significance was determined by paired two-tailed Student’s *t*-tests. **B.** Identification and enumeration of CD138^+^ thymic *i*NKT cells among BALB/c *Foxp3-GFP* reporter mice one week after injection of anti-DR3 or isotype control antibody (Ctrl IgG). The contour plot is representative, and the bar graphs are a summary of data from 11 independent experiments with a total of 11 mice for each group. Statistical significance was determined by paired two-tailed Student’s *t*-tests.

Because we identified NKT17 cells based on their CD138 expression (*Luo et al., 2021*), we could not exclude the possibility that DR3 ligation would appear to deplete NKT17 cells by downregulating CD138 expression. In fact, the shedding of the CD138 ectodomain is a well-described process that results in the loss of surface CD138 (*Rangarajan et al., 2020*), so that DR3 ligation might have triggered CD138 downregulation without altering the composition of the thymic *i*NKT cells. To determine whether DR3 ligation leads to the actual loss of NKT17 cells or if anti-DR3 only induces the downregulation of surface CD138 expression on NKT17 cells, we considered it necessary to identify NKT17 cells with markers other than surface CD138. Hence, we employed the surface markers CD4 and CD122 to discriminate individual *i*NKT subsets (*Georgiev et al., 2016*). CD122 is selectively expressed on NKT1 cells (*Won et al., 2021*), so that CD122^+^ *i*NKT cells correspond to the NKT1 subset. NKT2 cells are CD122-negative but they express large amounts of CD4 (CD122^−^CD4^+^). Most NKT17 cells, on the other hand, are negative for both CD4 and CD122 (*Georgiev et al., 2016*). In fact, CD122/CD4 double-negative (DN) cells were RORγt^+^ and expressed high levels of both DR3 and CD138, confirming that they corresponded to NKT17 cells (**Figure 3A**). Therefore, the combined use of CD122 and CD4 permitted us to identify NKT17 subset cells without using CD138. In agreement, DR3 was also highly expressed on the DN *i*NKT cells of CD138-deficient *Sdc1*^−/−^ mice, marking them as NKT17 cells, (**Figure 3B**). These results indicated that DR3 expression is a bona fide marker for thymic NKT17 cells, independently of CD138.

**Figure 3.**
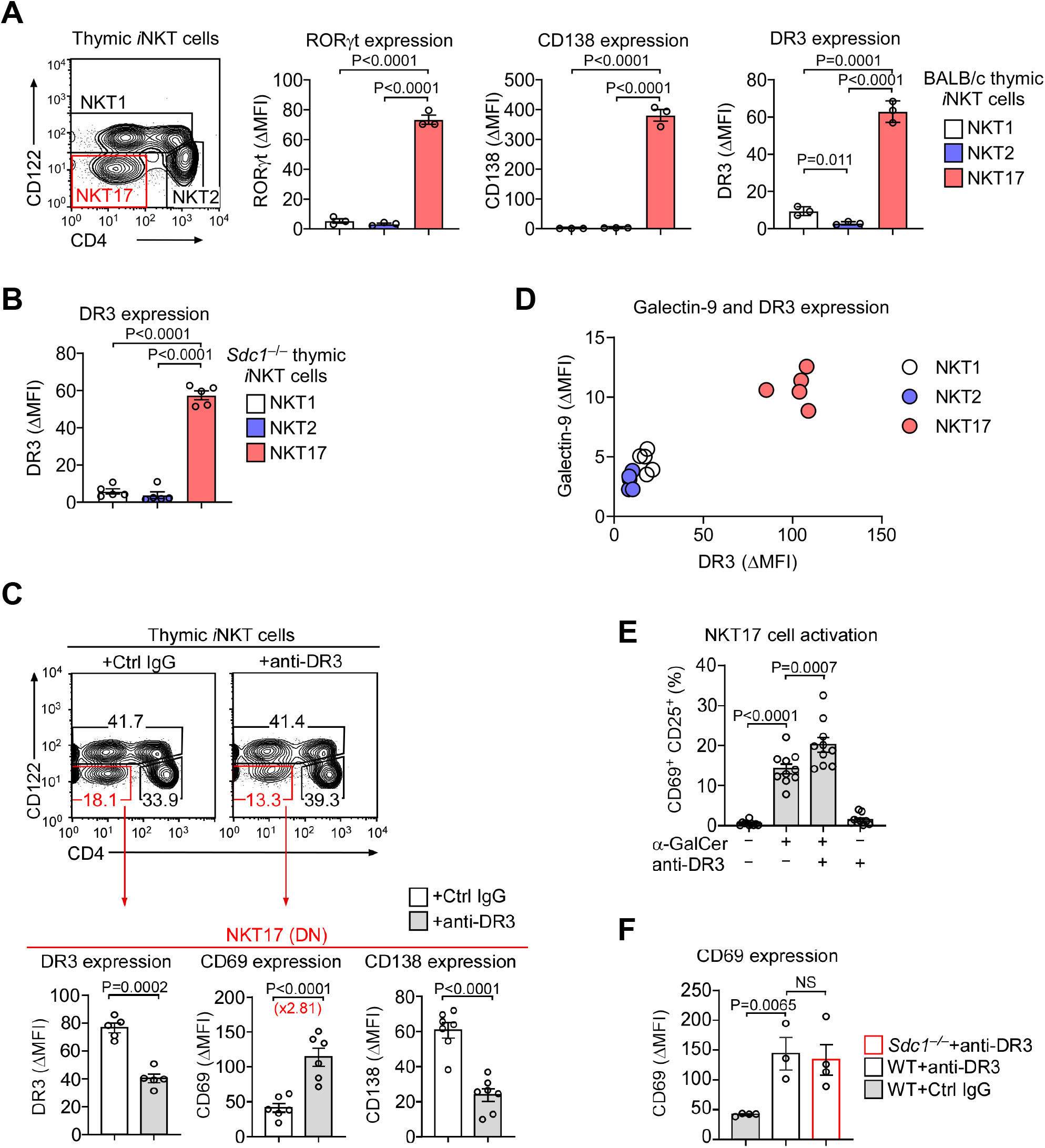
DR3 is a specific and functional marker of thymic NKT17 cells. **A.** Thymic *i*NKT subsets were identified by CD4 versus CD122 expression (contour plot), and the expression of subset-specific signature molecules were quantified for the indicated *i*NKT subsets (bar graphs). The contour plot is representative, and the bar graphs are summaries of data from three independent experiments with a total of 3 BALB/c mice. Statistical significance was determined by unpaired two-tailed Student’s *t*-tests. **B.** DR3 expression on CD4, CD122-disparate thymic *i*NKT subsets of CD138-deficient (*Sdc1*^−/−^) BALB/c mice. The bar graph shows the summary of data from three independent experiments with a total of 5 *Sdc1*^−/−^ BALB/c mice. Statistical significance was determined by unpaired two-tailed Student’s *t*-tests. **C.** Activation marker expression on thymic NKT17 cells of BALB/c *Foxp3*-GFP reporter mice upon anti-DR3 antibody injection. The contour plot is representative, and the bar graphs are summaries of data from 7 independent experiments with a total of 14 BALB/c *Foxp3-GFP* reporter mice injected either with anti-DR3 (7 mice) or isotype control antibodies (7 mice). Statistical significance was determined by paired two-tailed Student’s *t*-tests. **D.** DR3 and Galectin-9 co-expression on thymic *i*NKT subsets of BALB/c mice identified by the CD4 versus CD122 expression. The graph is a summary of data from 2 independent experiments with a total of 5 BALB/c mice. **E.** *In vitro* activation of thymic NKT17 cells by overnight stimulation with α-GalCer (100 ng/ml) in the presence or absence of anti-DR3 antibody (2 μg/ml) stimulation. The bar graph is a summary of data from 4 independent experiments with a total of 10 BALB/c mice. Statistical significance was determined by paired two-tailed Student’s *t*-tests. **F.** CD69 expression on thymic NKT17 cells from *Sdc1*^−/−^ and littermate control (WT) BALB/c mice one week after injection with anti-DR3 or isotype control antibodies (Ctrl IgG). The bar graph is a summary of data from 4 independent experiments with a total of 4 mice for each group. Statistical significance was determined by unpaired two-tailed Student’s *t*-tests. The following figure supplements are available for Figure 3: **Figure supplement 1**. CD69 expression upon DR3 injection in thymic NKT1 and NKT2 cells. **Figure supplement 2**. Galectin-9 and DR3 expression in thymic *i*NKT cell subsets of BALB/c mice. **Figure supplement 3**. *In vitro* stimulation of thymic *i*NKT cells with α-GalCer and/or anti-DR3 antibodies.

### DR3 ligation selectively activates thymic NKT17 cells

Equipped with this toolkit to identify NKT17 cells, we next assessed the effect of DR3 ligation on NKT17 cells. Injection of agonistic anti-DR3 antibodies into BALB/c mice induced the expression of CD69, a classical activation marker (*Ziegler et al., 1994*), on thymic NKT17 cells which was accompanied by decreased CD138 expression (**Figure 3C**). Consequently, the loss of CD138^+^ *i*NKT cells upon DR3 injection (**Figure 2B**) is unlikely due to the loss of NKT17 cells but more likely the result of their selective activation. Indeed, DR3-induced activation was largely limited to thymic NKT17 cells with minimal or no activation of NKT1 and NKT2 cells (**Figure 3C; Figure 3–figure supplement 1**). Importantly, DR3 signaling was reported to require the co-expression of galectin-9 (*Madireddi et al., 2017*), and we found that NKT17 cells were incidentally the only thymic *i*NKT subset that expressed both DR3 and galectin-9 (**Figure 3D; Figure 3–figure supplement 2**).

While the injection of anti-DR3 antibodies activated NKT17 cells *in vivo*, anti-DR3 antibodies alone were insufficient to induce their activation *in vitro* (**Figure 3E**). However, DR3 ligation significantly boosted the effect of α-GalCer stimulation and bolstered the expression of the activation markers CD25 and CD69 on NKT17 cells (**Figure 3E; Figure 3–figure supplement 3**), indicating that DR3 acts as a costimulatory molecule. Altogether, these results suggested that DR3 is a functional marker for NKT17 cells through which the *i*NKT immune response can be skewed towards IL-17 immunity. Finally, we examined whether CD138 is a prerequisite for DR3-induced activation of CD138 for NKT17 cells (*Dai et al., 2015*). Here, we found that *Sdc1*^−/−^ NKT17 cells still responded robustly to DR3 ligation so that the activation-induced upregulation of CD69 was comparable to that of WT NKT17 cells (**Figure 3F**). Therefore, CD138 is specifically expressed on NKT17 cells but not required for DR3-induced NKT17 activation.

Collectively, our results identified the cytokine receptor DR3 as a new costimulatory molecule that is specifically expressed on and activates thymic NKT17 cells. In this regard, DR3 represents a new class of immunomodulatory molecules whose expression and function are linked to a specific *i*NKT subset. These results open new avenues for elucidating how different *i*NKT subsets, that express the same invariant TCR and respond to the same agonistic glycolipid, *i.e*., α-GalCer, can elicit subset-specific immune responses *in vivo*.

## Materials and Methods

### Mice

BALB/cAnNCrl and C57BL/6 mice were purchased from the Charles River Laboratories. CD138-deficient (*Sdc1*^−/−^) mice and RORγt^Tg^ mice were previously described (*Alexander et al., 2000; Ligons et al., 2018; Luo et al., 2021*), and these animals were backcrossed in-house onto BALB/cAnNCrl background before analyses. Foxp3-DTR/EGFP mice were obtained from the Jackson Laboratory and maintained on BALB/cAnNCrl background (*Lahl et al., 2007*). Animal experiments were approved by the NCI Animal Care and Use Committee. All mice were cared for in accordance with the NIH guidelines.

### Antibodies

Antibodies specific for the following antigens were used for staining: TCRβ (H57-597), CD4 (GK1.5), CD24 (M1/69), CD138 (181-2), CD122 (TM-β1), CD44 (IM7), DR3 (4C12), Galectin-9 (108A2), CD69 (H1.2F3), CD25 (PC61.5), IL-7Rα (A7R34), IL-17 (eBio17B7), PLZF (9E12), and RORγt (Q31-378). Armenian Hamster IgG isotype Control Antibody (HTK888) was used as control for anti-DR3 staining. Rat IgG2a, κ Isotype Ctrl (RTK2758) was used as control for anti-Galectin-9 staining. PBS-57-loaded mouse CD1d tetramers were obtained from NIH Tetramer Core Facility (Emory University, Atlanta, GA).

### Enrichment of mature thymocytes

CD24-negative mature thymocytes were enriched by magnetic depletion of CD24^+^ cells, as previously described (*Park,Kwon, et al., 2019*). In brief, total thymocytes were processed to single cell suspension in 10% FBS/HBSS (20×10^6^ cells/ml) and incubated with rat anti-mouse CD24 antibodies (M1/69, Biolegend) (30 μg/100 ×10^6^ cells) for 30 mins on ice. After washing off excess reagents, thymocytes were mixed with anti-rat IgG-conjugated BioMag beads (QIAgen) and incubated for 45 mins at 4°C on a MACSmix Tube Rotator (Miltenyi Biotec). Anti-CD24 antibody-bound cells were then magnetically removed, and non-binding cells were harvested for further experiments.

### Flow cytometry

Fluorescence antibody-stained single-cell suspensions were analyzed using LSRFortessa or LSRII flow cytometers (BD Biosciences). For live cell analysis, dead cells were excluded by adding propidium iodide before running the samples on flow cytometers. For fixed cell staining and analysis, cells were stained with Ghost Dye Violet 510 (Tonbo) for exclusion of dead cells, followed by surface staining and fixation with Foxp3 fixation buffer (eBioscience). Afterwards, cells were permeabilized using reagents from the Foxp3 intracellular kit according to the manufacturer’s instructions (eBioscience). Excess reagents were removed by extensive washing in FACS buffer (0.5% BSA, 0.1% sodium azide in HBSS) before analysis.

### Identification of iNKT subsets by intracellular staining

Thymic *i*NKT subsets were identified by staining for transcription factors as previously described (*Park,DiPalma, et al., 2019*). In brief, thymocytes were stained with fluorescence-conjugated PBS-57-loaded mouse CD1d tetramers, followed by antibody staining for other surface markers for 40 minutes. After washing out excess reagents, cells were fixed in 150 μl of a 1:3 mixture of concentrate/diluent working solution of the Foxp3 Fixation Buffer and further diluted with 100 μl FACS buffer. After 20 minutes at room temperature, cells were washed twice with permeabilization buffer (eBioscience) before adding antibodies for transcription factor staining. After 1 hour of incubation at room temperature, cells were washed, resuspended in FACS buffer, and analyzed by flow cytometry.

### Anti-DR3 agonistic antibody injection

For *in vivo* anti-DR3 ligation, mice were injected i.p. with either 10 μg anti-DR3 antibody (4C12, Biolegend) or 10 μg Armenian Hamster IgG control antibody (HTK888, Biolegend). One week after injection, thymus and spleen were harvested for further analysis.

### *In vitro* stimulation of thymic *i*NKT cells

Single cell suspension of freshly isolated thymocytes were plated into 24-well plates at 2×10^6^ cells/mL with 100 ng/mL of α-GalCer in the presence or absence of anti-DR3 antibody (2 μg/mL) or with anti-DR3 antibody alone (10 μg/mL) (*Schreiber et al., 2010*). Cells were cultured overnight at 37°C in a 7.5% CO_2_ incubator before analysis by flow cytometry.

### Statistics

Data are shown as the mean ± SEM. Two-tailed Student’s *t*-test was used to calculate P values. P values of less than 0.05 were considered significant, where NS indicates not significant. Statistical data were analyzed using the GraphPad Prism 8 software.

## Acknowledgement

This study was supported by the Intramural Research Program of the US National Institutes of Health, National Cancer Institute, Center for Cancer Research.

## Authorship Contribution

SL and NL designed and performed the experiments, analyzed the data, and contributed to the writing of the manuscript. AC performed experiments, analyzed the data, and commented on the manuscript. JP conceived the project, analyzed the data, and wrote the manuscript.

---

**Figure 1–figure supplement 1.**
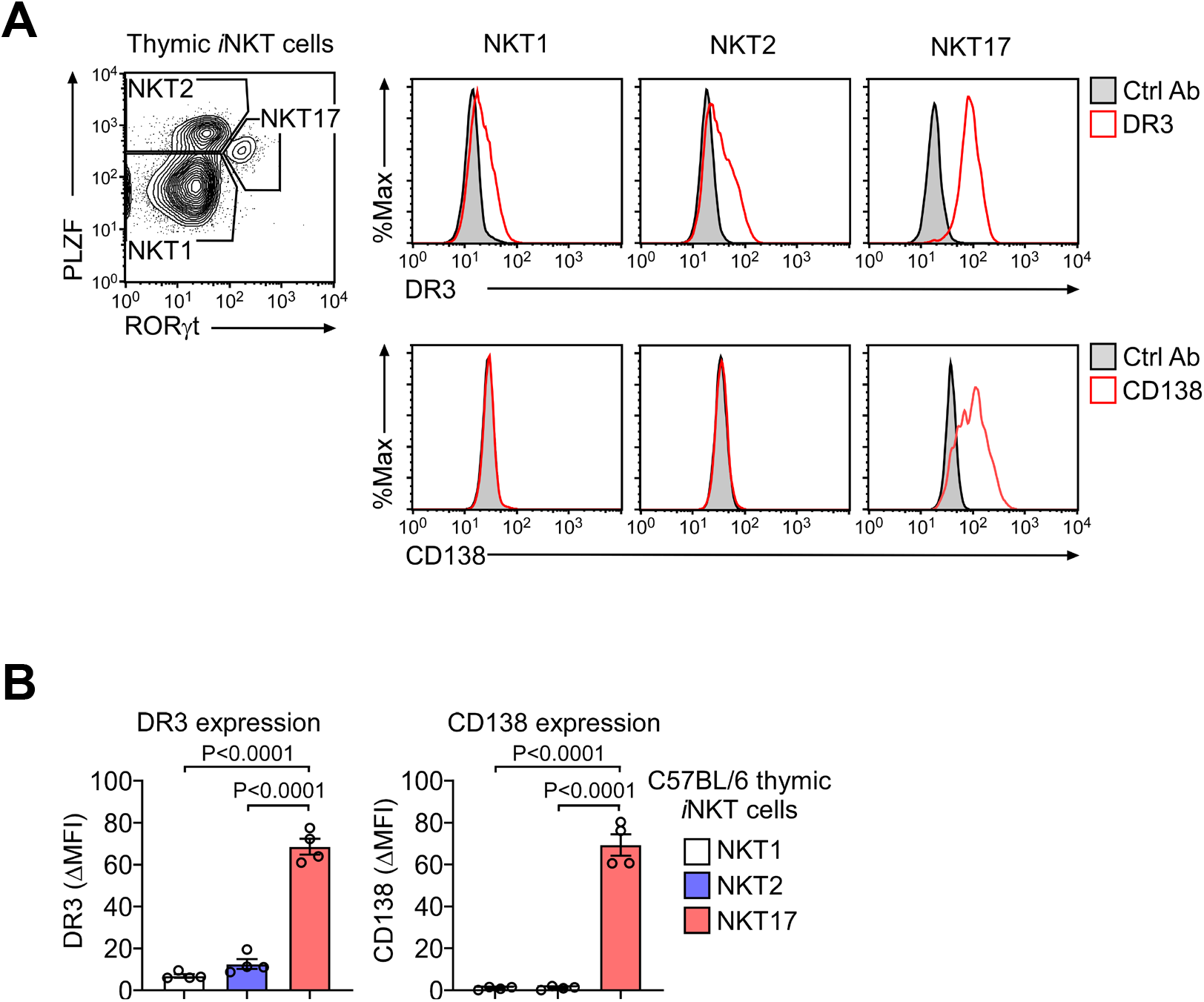
DR3 expression on thymic NKT17 cells of C57BL/6 mice. **A**. *i*NKT subsets were identified among thymocytes of C57BL/6 mice by intracellular staining for RORγt and PLZF and assessed for subset-specific expression of DR3 and CD138. The data are representative of 3 independent experiments. **B**. Bar graphs show DR3 expression (ΔMFI) (left) and CD138 expression (ΔMFI) (right) among thymic subsets of C57BL/6 mice as identified by intracellular staining for RORγt and PLZF. The data are from 3 independent experiments with a total of 4 pooled C57BL/6 mice and presented as mean ± SEM. Statistical significance was determined by unpaired two-tailed Student’s *t*-tests.

**Figure 1–figure supplement 2.**
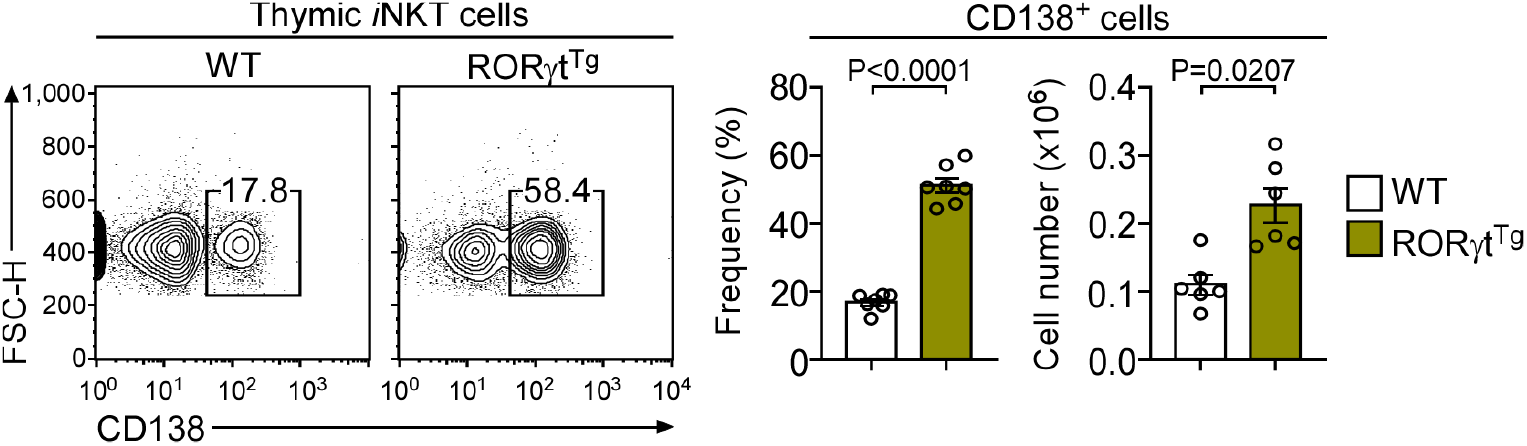
Thymic CD138^+^ *i*NKT cells in WT and RORγt^Tg^ BALB/c mice. Contour plots show CD138 versus FSC-H of *i*NKT cells in littermate control (WT) BALB/c and RORγt^Tg^ BALB/c mice (left). Bar graphs show the frequency and cell number of CD138^+^ *i*NKT cells in WT BALB/c and RORγt^Tg^ BALB/c mice (right). Contour plots are representative and bar graphs show the summary of 3 independent experiments with a total of 6 WT and 6 RORγt^Tg^ BALB/c mice. Data are presented as mean ± SEM. Statistical significance was determined by paired two-tailed Student’s *t*-tests.

**Figure 3–figure supplement 1.**
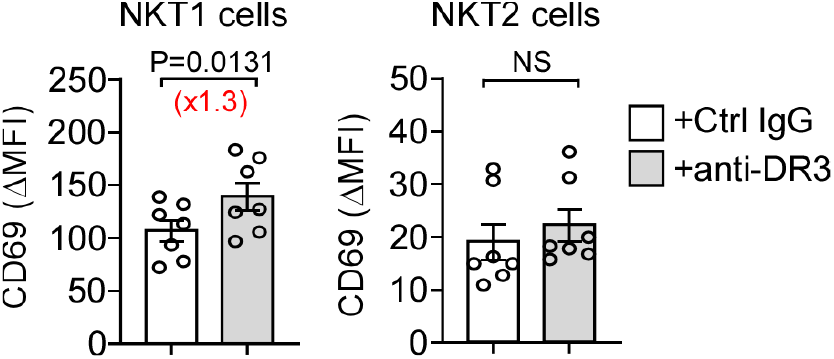
CD69 expression upon DR3 injection in thymic NKT1 and NKT2 cells. Bar graphs show the CD69 expression (ΔMFI) of thymic NKT1 (left) and NKT2 (right) cells in Foxp3-DTR/EGFP BALB/c mice, one week after injection with anti-DR3 or isotype control antibodies. The results are summary of 7 independent experiments with a total of 14 mice injected with either anti-DR3 antibodies (7 mice) or with isotype control antibodies (7 mice). Data are presented as mean ± SEM. NS, non-significant. Statistical significance was determined by paired two-tailed Student’s *t*-tests.

**Figure 3–figure supplement 2.**
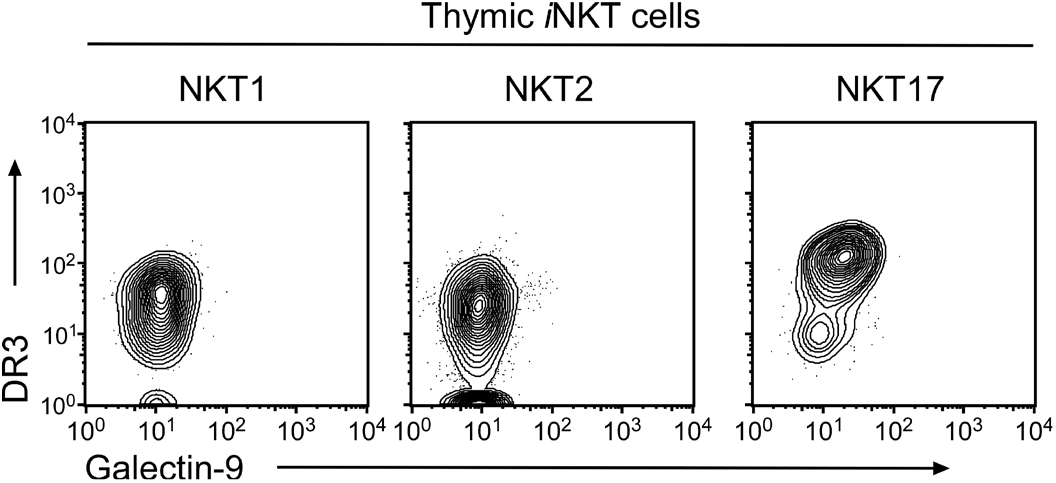
Galectin-9 and DR3 expression in thymic *i*NKT cell subsets of BALB/c mice. The counter plots show galectin-9 versus DR3 profiles of thymic NKT1 cells (CD122^+^), NKT2 cells (CD122^−^CD4^+^) and NKT17 cells (CD122^−^CD4^−^). The data are representative of 2 independent experiments with a total of 5 WT BALB/c mice.

**Figure 3–figure supplement 3.**
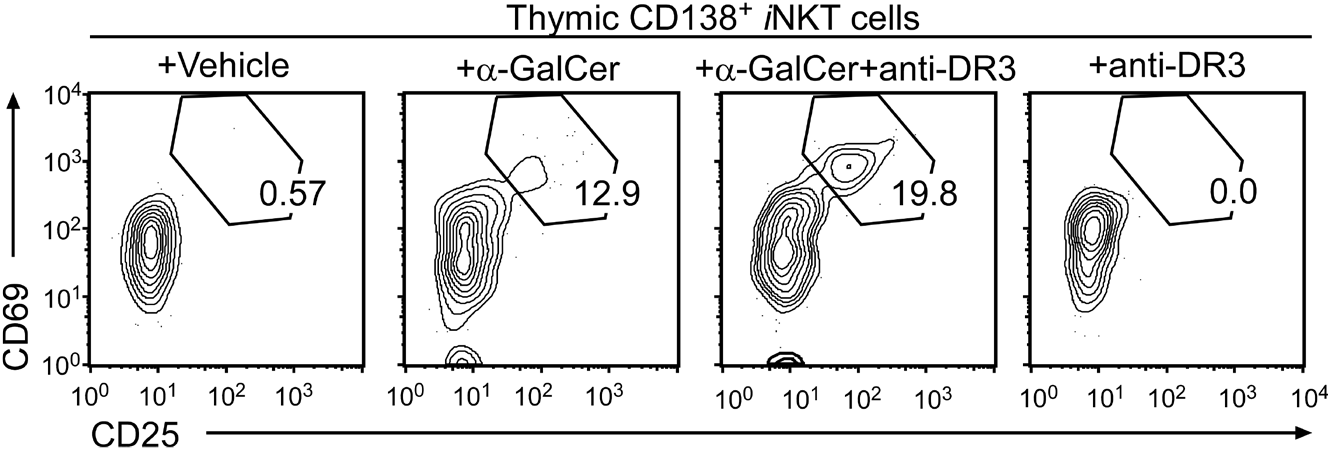
*In vitro* stimulation of thymic *i*NKT cells with α-GalCer and/or anti-DR3 antibodies. Contour plots show CD69 versus CD25 profiles of CD138^+^ thymic *i*NKT cells of BALB/c mice that were cultured O/N with α-GalCer and/or anti-DR3 antibodies. The data are representative of 4 independent experiments with a total of 10 WT BALB/c mice for each group.

